# Differentiation genes were governed by DNA methylation during hair follicle morphogenesis in Cashmere goat

**DOI:** 10.1101/2020.01.30.926360

**Authors:** Shanhe Wang, Fang Li, Jinwang Liu, Yuelang Zhang, Yujie Zheng, Wei Ge, Lei Qu, Xin Wang

**Author notes:** These authors contributed equally to this work (Shanhe Wang, Fang Li). Correspondence (L.Q.) and (X.W.).

## Abstract

DNA methylation plays a critical role in early embryonic skin development by controlling gene expression. Act as an indirect regulator, long non-coding RNA (lncRNA) recruit DNA methyltransferases to specific genomic sites to methylate DNA. However, the molecular regulation mechanisms underlying hair follicle morphogenesis is unclear in cashmere goat. In this study, RNA-seq and Whole-genome bisulfite sequencing (WGBS) in embryonic day 65 (E65) and E120 skin tissues of cashmere goat were used to reveal this complex regulatory process. RNA-seq, qRT-PCR and immunohistochemistry results showed that Wnt signaling played an important role in both hair follicle induction and differentiation stage, transcriptional factors (TFs) including Hoxc13, Sox9, Sox21, Junb, Lhx2, Vdr and Gata3 participated in hair follicle differentiation via specific expression at E120. Subsequently, combination of WGBS and RNA-seq analysis showed that the expression of hair follicle differentiation genes and TFs genes was negatively correlated with DNA methylation level generally. A portion of hair follicle differentiation genes were methylated and repressed in hair follicle induction stage but were subsequently demethylated and expressed during hair follicle differentiation stage, suggesting DNA methylation play an important role in hair morphogenesis through regulating associated gene expression. Furthermore, the potential differentially expressed lncRNAs associated with DNA methylation on target gene were revealed. LncRNA XR_001918556 may affect the DNA methylation of TFs gene *Gata3*, lnc-003786 may affect the DNA methylation of signaling gene *Fgfr2*. In conclusion, differentiation genes were governed by DNA methylation, resulting in repressed expression in hair follicle induction stage and high expression in hair follicle differentiation stage. Furtherly, potential lncRNAs associated with DNA methylation on target genes were delineated. This study would enrich the regulatory network and molecular mechanisms on hair morphogenesis.

## Introduction

Hair is a primary characteristic of mammals, and exerts a wide range of functions including thermoregulation, physical protection, sensory activity, and social interactions (Paus and Cotsarelis, 1999; Schneider et al., 2009). Cashmere is upmarket textile material produced by the secondary hair follicle with high economic values (Ge et al., 2018; Wang et al., 2017). As the number and quality of cashmere depend on cashmere morphogenesis, it is therefore of great value to dissect the critical genes, signaling pathways and their regulatory machinery underlying hair follicle morphogenesis in cashmere goat.

Hair follicle morphogenesis takes place during embryonic skin development, which relies on tightly coordinated ectodermal–mesodermal interactions (Biggs and Mikkola, 2014; Botchkarev and Kishimoto, 2003; Millar, 2002; Schmidt-Ullrich and Paus, 2005). Researches in mice showed that hair follicle morphogenesis is initiated after secreted epidermal Wnts activating broad dermal Wnt signaling (Chen et al., 2012), which in turn through unknown dermal signaling and subsequent Wnt, Eda and FGF20 epidermal downstream signaling, leads to hair placode (Pc) induction in epidermis (Lee and Tumbar, 2012; Schneider et al., 2009; Wang et al., 2012)and dermal condensate (DC) formation below (Huh et al., 2013; Mok et al., 2019). Following the induction stage, hair follicle enter organogenesis and subsequent cytodifferentiation stage, in which Pc cells give rise to all the epithelial components of full-developed hair follicle including outer root sheath, inner root sheath, hair matrix, hair shaft and hair follicle stem cell, while the DC cells will develop into the follicular dermal papilla and connective tissue sheath (Asakawa et al., 2017; Avigad Laron et al., 2018; Mesler et al., 2017). A number of molecules and their interactions in each phase that play a role in hair follicle development have been identified using transgenic mice model and hair follicle regeneration assay (Bak et al., 2018; Glover et al., 2017; Nakamura et al., 2013). However, the unique molecular features of specific cell type and the regulatory relationships between signaling pathways involved in these processes are largely unknown(Sennett et al., 2015), especially in cashmere.

Hair follicle morphogenesis results from the process of temporal-spatial expression of genes under the control of genetic and epigenetics, while DNA methylation has been shown to be implicated in the regulation of cell-or tissue-specific gene expression during embryogenesis (Michael et al., 2007; Reik and Dean, 2001). DNA methylation undergoes dynamic remodeling during early embryogenesis to initially establish a globally demethylated state and then subsequently, a progressively lineage-specific methylome that maintains cellular identity and genomic stability (Baubec and Schubeler, 2014; Senner, 2011). As development and differentiation proceed, differentiated cells accumulate epigenetic marks that differ from those of pluripotent cells, and differentiated cells of different lineages also accumulate different marks (Bock et al., 2012; Feng et al., 2010; Suzuki and Bird, 2008). However, it is still unknown about the function of DNA methylation in regulating cell lineage specification during hair morphogenesis.

DNA methyltransferases (DNMTs) involved in DNA methylation lack sequence-specific DNA binding motifs, while many lncRNAs have DNA- and protein-binding motifs, allowing them to carry DNMTs to specific genomic sites (Carlson et al., 2015; Mohammad et al., 2012). Emerging data indicate that lncRNAs function as guides and tethers, and may be the molecules of choice for epigenetic regulation (Chen et al., 2019). Meanwhile, previous studies revealed that lncRNA regulates hair follicle stem cell proliferation and differentiation (Cai et al., 2018). However, whether lncRNA mediate DNA methylation and contribute to hair morphogenesis in cashmere goat is unknown.

To investigate the molecular identity and regulatory mechanism underlying hair morphogenesis in cashmere goat, RNA-seq was conducted on skin samples at hair follicle induction and differentiation stages from E 65 and E 120, we composed a molecular snapshot of an entire tissue, and uncovered genes in cell-type-specific signatures through transcriptome cross-comparisons with mice. Furthermore, genome-wide DNA methylation profiles between skin tissues in E 65 and E 120 were investigated using WGBS. Through integrated analysis of mRNA and lncRNA transcriptome with WGBS data, the regulation of DNA methylation on hair induction and differentiation and the potential lncRNAs involved in DNA methylation to take part in hair morphogenesis have been delineated. Our work would enrich the underlying molecular mechanisms of hair follicle morphogenesis and skin development.

## Materials and Methods

### Animals

Shanbei White Cashmere goats with fine fiber production trait were used in this study. All the goats were obtained from Shanbei cashmere goats engineering technology research center of Shaanxi province, China. The experimental animals were fed according to the local cashmere goat standard of Shaanxi (DB61/T583-2013, http://www.sxny.gov.cn/). According previous morphology studies on hair morphogenesis of cashmere goat, in which hair follicle induction initiated around E 65 and hair follicle differentiation thrived around E 120, six pregnant Shanbei White Cashmere goats (two years old, weighing 30 – 40 kg) were selected to obtain fetal skin samples at E 65 and E 120 (n=3). Each time point had three replicates. After intravenous injection of Rompun (0.3 mg/kg) to anesthesia, six fetuses were delivered from six different females by caesarean operation. Skin samples were collected from the right mid-side of the fetuses, rinsed in ice-cold DEPC-treated water and cut into small pieces. At the same time, other tissues including muscle, adipose, heart, liver, spleen, lungs, kidney, duodenum and gonad were collected. Every tissue sample was divided into two parts; one was fixed with 4 % paraformaldehyde and another one was frozen in sample protector for RNA/DNA (Takara, China) and stored at −80 °C for subsequent analysis. The carcasses were frozen to designated location waiting bio-safety disposal.

All the experimental procedures with goats used in the present study had been given prior approval by the Experimental Animal Manage Committee of Northwest A&F University (2011-31101684). All the operations and experimental procedures were complied with the national standard of Laboratory Animal-Guideline for Ethical Review of Animal Welfare (GB/T 35892-2018) and Guide for the Care and Use of Laboratory Animals: Eighth Edition (Council, 2011).

### Transcriptome sequencing and bioinformatics analysis

To obtain a transcriptome reference between E 65 and E 120, total RNA was extracted from the collected skin and other tissues. The RNA concentration and quality were determined using the Agilent 2100 Bioanalyzer (Agilent Technologies, USA). RNA-seq was performed as previously described (Li et al., 2018). We used the skin RNA samples to construct RNA-seq libraries from E 65 and E 120. Clean data were obtained by trimming reads containing adapter, reads containing over 10 % of ploy-N, and low-quality reads (> 50 % of bases whose Phred scores were < 20) from the raw data. Then all subsequent analysis was based on the high-quality data. Then, the high quality reads were mapped to the goat genome v2.0 (ftp://ftp.ncbi.nlm.nih.gov/genomes/all/GCA/000/317/765/GCA_000317765.2_CHIR_2.0) using Bowtie v2.0.6 (Langmead et al., 2009) and the mapped reads for each sample were assembled using Cufflinks (v2.2.1). The differential expression changes were calculated for the pairwise comparison between E 65 and E 120 skin tissues, transcripts or genes with a P-adjust ≤ 0.05 (Storey, 2003) and fold change ≥ 2 were described as differentially expressed. To explore the function of lncRNAs, we predicted the target genes of lncRNAs in *cis* and *trans*. And pearson’s correlation coefficients were calculated between expression levels of lncRNAs and mRNAs with custom scripts (Pearson correlation ≥ 0.95 or ≤ −0.95).

Gene Ontology (GO) enrichment analysis of differentially expressed genes was implemented using Gene Ontology Consortium (http://www.geneontology.org/) (Chibucos, 2015). Gene ontology terms with corrected *P* value less than 0.05 were considered significantly enriched by differentially expressed genes. Pathway analysis was used to identify significant pathways for the differentially expressed genes according to the Kyoto Encyclopedia of Genes and Genomes (KEGG) (http://www.genome.jp/kegg/) (Kanehisa et al., 2008). We used KOBAS software (main parameter: blastx 1e-10; padjust: BH) to test the statistical enrichment of differentially expressed genes in KEGG pathways (Mao et al., 2005).

### Quantitative Real-time PCR (qRT-PCR)

The first-strand cDNA was obtained using a PrimeScript™ RT reagent Kit with gDNA Eraser (TAKARA, China), and then were subjected to quantification of the mRNAs with β-actin as an endogenous control on the Bio-Rad CFX96 Touch™ Real Time PCR Detection System (Bio-Rad, USA). The qRT-PCR reaction consisted of 10 μL 2 ×SYBR^®^ *Premix Ex Taq™* II (TAKARA, China), 0.8 μL specific forward/reverse primer (10μM), 1 μL cDNA, and ddH_2_O to a final volume of 20 μL. The qRT-PCR was performed using the following conditions: 95 °C for 60 s, 40 cycles of 95 °C for 10 s, and the optimized annealing temperature for 30 s. Semi-quantitative RT-PCR was performed on 2720 thermal cycler (Applies biosystems) machine using ES Taq master mix (Cwbio, China). Primers used were provided in Table S1.

Differences between samples at E 65 and E 120 (n=3) were calculated based on the 2^-ΔΔCt^ method and normalized to β-actin. Measurements were recorded in duplicate. Differences in gene expression between the groups were detected by independent sample *t*-test.

### Histology and immunohistochemistry (IHC)

Skin samples from E65 and E120 were fixed with 4 % paraformaldehyde, followed by dehydration further embedded in paraffin and cut into 5 μm sections with a microtome (Leica RM2255, Nussloch, Germany). Sections were rehydrated, blocked with 10 % goat serum and 3 % bovine serum albumin (Sigma, USA), and incubated 40 min at room temperature. Primary antibody against interest protein was then incubated with the samples at 4 °C overnight. Primary antibodies used were: Bmp2 (Abcam, Cat. No. ab214821, rabbit 1:200), Sox9 (Abcam, Cat. No. ab185966, rabbit 1:200), Vdr (Proteintech, Cat. No. 14526-1-AP, rabbit 1:150), Sox2 (Proteintech, Cat. No. 11064-1-AP, rabbit 1:150), Bmp4 (Proteintech, Cat. No. 12492-1-AP, rabbit 1:150), β-catenin (Proteintech, Cat. No. 51067-2-AP, rabbit 1:150), Wls (Proteintech, Cat. No. 17950-1-AP, rabbit 1:100), Fzd10 (Proteintech, Cat. No. 18175-1-AP, rabbit 1:150), Edar (Sangon Biotech, Cat. No. D160287, rabbit 1:100), Fgf20 (Sangon Biotech, D161681, rabbit 1:100). Subsequently, fluorescent goat anti-rabbit Ig-CY3/FITC-conjugated secondary antibody (Beyotime biotechnology, Cat. No. A0516/A0562, goat, 1:100) or HRP-conjugated secondary antibody (Sangon Biotech, Cat. No. 110058, goat, 1:100) were used to specifically bind to primary antibody. Metal Enhanced DAB Substrate Kit (Solarbio, China) was used to color developing under the catalysis of HRP. Hoechst33342 (Beyotime biotechnology, China) was used for nuclei staining and the slides were finally mounted with Vecatshield mounting media (Vector, USA). H&E staining were performed according to standard procedures. Fluorescent pictures were taken under LEICA TCS SP5 II confocal microscopy (Leica Microsystems GmbH, Wetzlar, Germany). All images of H&E stained sections were taken on an Eclipse 80i microscope (Nikon, Japan).

### DNA extraction, WGBS library construction and sequencing

Genomic DNA was extracted from skin samples (E 65 and E 120) using Qiagen DNeasy Blood & Tissue Kit (Qiagen, USA) according to the manufacturer’s instructions. Genomic DNA degradation and contamination was monitored on agarose gels. DNA purity and concentration were checked using the NanoPhotometer^®^ spectrophotometer (IMPLEN, USA).

WGBS was performed as previously described (Li et al., 2018) in E65 and E120 skin tissues (n=3) of cashmere goat. A total of 5.2 μg of genomic DNA spiked with 26 ng lambda DNA was fragmented by sonication to 200 – 300 bp with Covaris S220, followed by end repair and adenylation. Cytosine-methylated barcodes were ligated to sonicated DNA according manufacturer’s instructions. Then these DNA fragments were treated twice with bisulfite using EZ DNA Methylation – GoldTM Kit (Zymo Research, USA), before the resulting single – strand DNA fragments were PCR amplificated using KAPA HiFi HotStart Uracil + ReadyMix (2X). Library concentration was quantified by Qubit^®^ 2.0 Flurometer (Life Technologies, CA, USA) and quantitative PCR, and the insert size was assayed on Agilent Bioanalyzer 2100 system.

The libraries were sequenced on an Illumina Hiseq 4000 platform and 150 bp paired-end reads were generated. Image analysis and base calling were performed with Illumina CASAVA pipeline. We use FastQC (fastqc_v0.11.5) to perform basic statistics on the quality of the raw reads. Then, those reads sequences produced by the Illumina pipleline in FASTQ format were pre-processed through Trimmomatic (Trimmomatic-0.36) software using the parameter (SLIDINGWINDOW: 4:15; LEADING:3, TRAILING:3; ILLUMINACLIP: adapter.fa: 2: 30: 10; MINLEN:36). The remaining reads that passed all the filtering steps was counted as clean reads and all subsequent analyses were based on this.

### Date analysis, identification of DMRs and functional enrichment analysis

For reads mapping to the reference genome, Bismark software (version 0.16.3) (Krueger and Andrews, 2011)was used to perform alignments of bisulfite-treated reads to reference genome (-X 700 --dovetail). The reference genome was firstly transformed into bisulfite-converted version (C-to-T and G-to-A converted) and then indexed using bowtie 2 (Langmead and Salzberg, 2012). Sequence reads were also transformed into fully bisulfite-converted versions (C-to-T and G-to-A converted) before they were aligned to similarly converted versions of the genome in a directional manner. Sequence reads that produce a unique best alignment from the two alignment processes (original top and bottom strand) were then compared to the normal genomic sequence and the methylation state of all cytosine positions in the read was inferred. The same reads that aligned to the same regions of genome were regarded as duplicated ones. The sequencing depth and coverage were summarized using deduplicated reads. The results of methylation extractor (bismark_methylation_extractor, -- no_overlap) were transformed into bigWig format for visualization using IGV browser. The sodium bisulfite non-coversion rate was calculated as the percentage of cytosine sequenced at cytosine reference positions in the lambda genome.

To identify the methylation site, we modeled the sum Mc of methylated counts as a binomial (Bin) random variable with methylation rate:

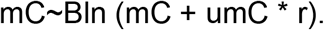

In order to calculate the methylation level of the sequence, we divided the sequence into multiple bins within 10 kb in size. The sum of methylated and unmethylated read counts in each window were calculated. Methylation level (ML) for each window or C site shows the fraction of methylated Cs, and is defined as:

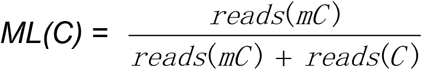

Calculated ML was further corrected with the bisulfite non-conversion rate according to previous studies (Ryan et al., 2013). Given the bisulfite non-conversion rate r, the corrected ML was estimated as:

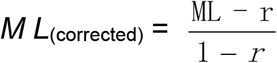

For differentially methylated analysis between the two age groups, differentially methylated regions (DMRs) were identified using the DSS software (Hao et al., 2014; Yongseok and Wu, 2016). According to the distribution of DMRs through the genome, we defined the genes related to DMRs as genes whose gene body region (from TSS to TES) or promoter region (upstream 2 kb from the TSS) have an overlap with the DMRs. GO enrichment and KEGG pathway analyses were conducted for the differentially methylated and expressed genes to investigate their biological processes and functions.

### Bisulfite sequencing polymerase chain reaction (BSP-PCR)

BSP-PCR was performed as we previously described (Wang et al., 2018) using E 65 and E 120 skin tissues genomic DNA. Every stage included three biological repetition. DNA treatment with sodium bisulfite was performed using the EZ DNA Methylation Kit (Zymo Research, USA) according to the manufacturer’s protocol, except that the conversion temperature was changed to 55 °C. The modified DNA samples were diluted in 10 μL of distilled water and should be immediately used in BSP or stored at – 80 °C until PCR amplification. The BSP primers were designed by the online MethPrimer software (http://www.urogene.org/methprimer/). The sequences of PCR primers used for amplifying the targeted products were shown Table S1. We used hot start DNA polymerase (Zymo Taq^TM^ Premix, Zymo Research, USA) for BSP production. PCR was performed in 50 μL of reaction volume, containing 200 ng genomic DNA, 0.3 μM of each primer, Zymo Taq^TM^ Premix 25 μL. The PCR was performed with a DNA Engine Thermal Cycler (Bio-Rad, USA) using the following program: 10 min at 95 °C, followed by 45 cycles of denaturation for 30 s at 94 °C, annealing for 40 s at 52 °C and extension for 30 s at 72 °C, with a final extension at 72 °C for 7 min. The PCR products were gel purified using Gel Purification Kit (Sangon, China), and then subcloned into the pGEM T-easy vector (Promega, USA). Different positive clones for each subject were randomly selected for sequencing (Sangon, China). We sequenced at least 5 clones from each independent set of amplification and cloning, hence, there were more than 15 clones for each DMR at E 65 and E 120 stage. The final sequence results were processed by online QUMA software 15 (http://quma.cdb.riken.jp/top/index.html).

## Results

### The morphology of hair follicle induction and differentiation stages in cashmere goat

Firstly, the corresponding hair follicle morphogenetic stages form E 65 and E 120 fetus cashmere skin were identified by H&E staining. As revealed by the H&E staining assay, hair follicle morphogenesis of cashmere goat initiated around E 65 with the characteristics of crowded epidermal keratinocytes, which showed enlarged and elongated, and got organized in microscopically recognizable hair placode (Pc). Meanwhile, Pc formation was succeeded by along with dermal condensate (DC) of specialized fibroblasts in the underlying mesenchyme (**Fig. 1a** and **c**). Up to E 120, the majority of primary hair follicles had matured with complete structure and hair shaft had emerged through epidermis, while the hair canal of secondary hair follicle was visible and the hair shaft began to grow up (**Fig. 1b** and **d**). In general, E 65 represented the induction stage, while the E 120 represented the differentiation stage of hair follicle morphogenesis.

**Fig 1.**
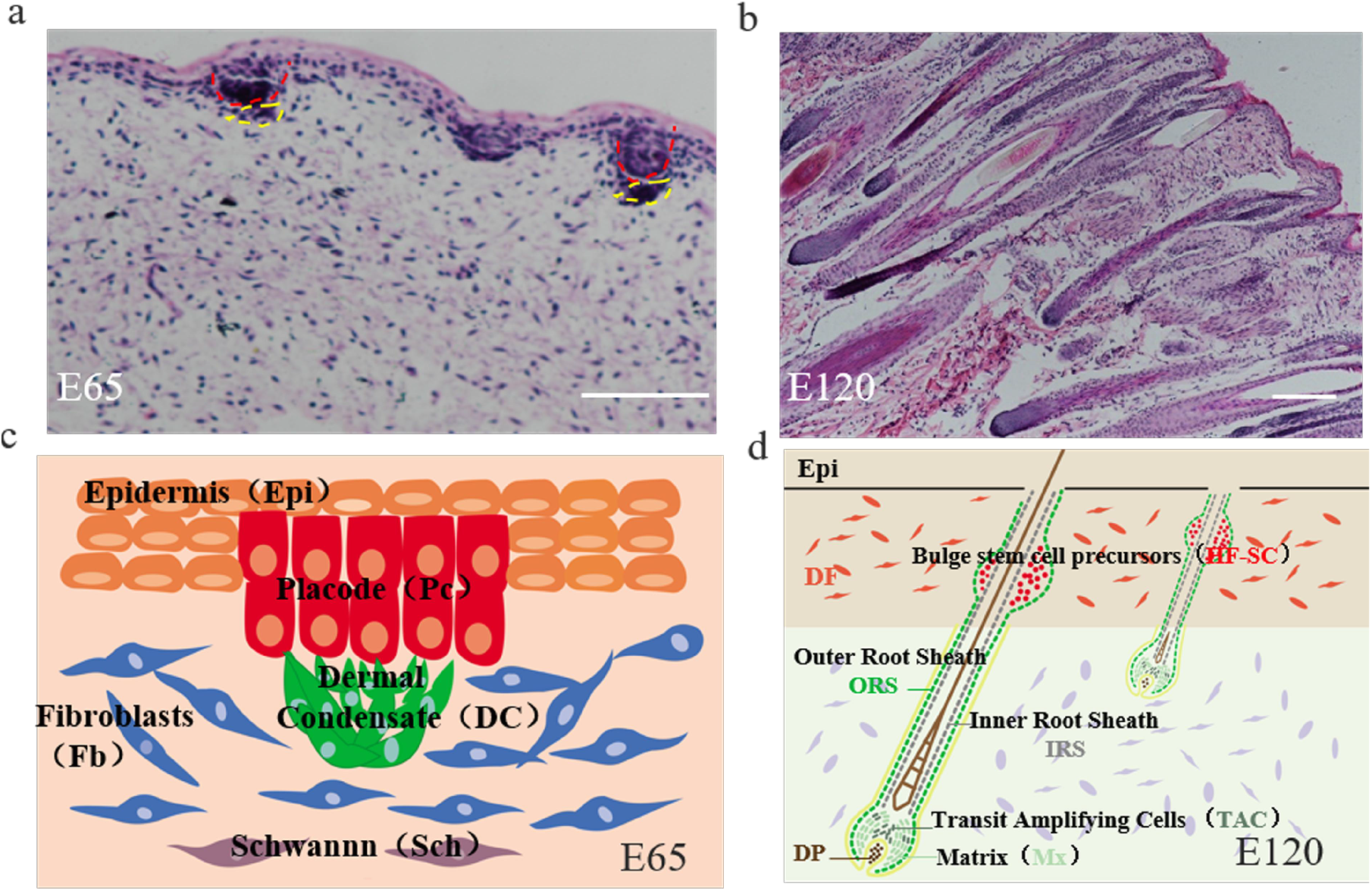
The skin morphology of E65 and E120 during hair morphogenesis in Shanbei White Cashmere goat. (**a-b**) The skin morphology of E65 and E120 during hair morphogenesis detected by H&E staining (Scale bars, 50 μm); (**c-d**) Schematic diagram of the skin morphology in E65 and E120 in Shanbei White Cashmere goat. Red dashed lines indicate epidermal hair follicle placode, Yellow dashed lines indicate dermal condensate.

### Defining distinct molecular signatures of hair follicle induction and differentiation

To reveal the distinct molecular signatures underlying hair follicle induction and differentiation in cashmere goat, we performed RNA-seq on E 65 and E 120 skin tissues using Illumina Hiseq 4000 system (n=3) (**Fig. 2a**). This approach resulted in a high-quality output, about 94.9 % index reads with quality score (Q score) >30 for all samples. On average, 99 million total clean reads and 93 million aligned reads were produced per sample. Next, we proceeded by mapping, aligning, and quantifying these reads to compute differentially expressed genes between E65 and E120 stages.

**Fig 2.**
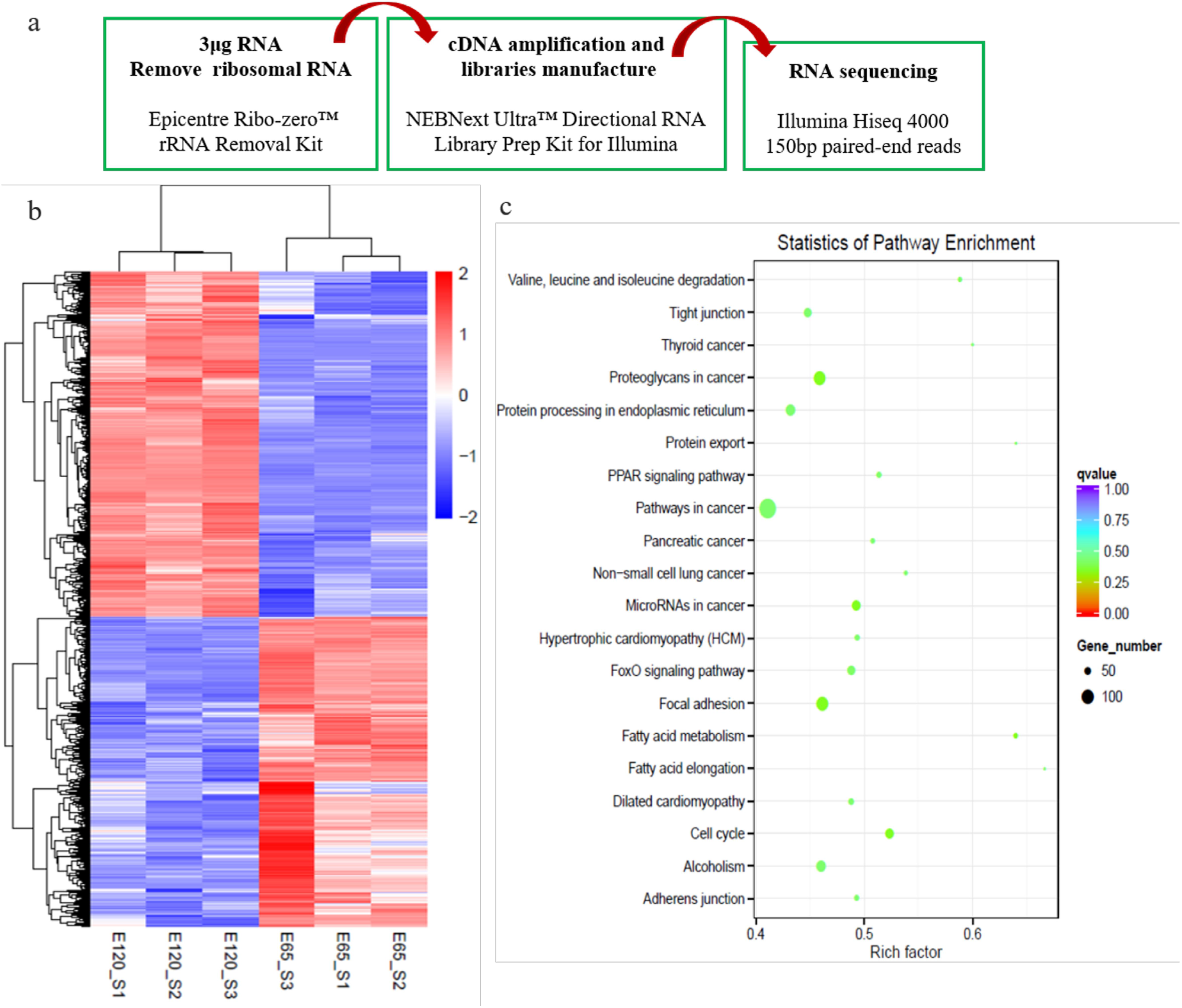
Critical signals and genes for hair follicle induction and differentiation stages revealed by RNA-sequencing and subsequent molecular verification. (**a**) Workflow of sample preparation for RNA sequencing. (**b**) The heatmap of DEGs between E65 and E120. (**c**) KEGG analysis of DEGs between E65 and E120.

Through comparing the RNA-seq data between E 65 and E 120, a total of 3,666 differential expressed genes (DEGs, Fold Change ≥ 2 and P-adjust value ≤ 0.05) were found, in which 1,729 genes were down regulated and 1,937 genes were up regulated in E 120 compared with E 65 (**Fig. 2b**) (Additional file 1). KEGG analysis of the DEGs revealed significant functional enrichment of cell migration and aggregation, highlighting the central roles of intercellular crosstalk and dynamic cell rearrangement in promoting skin and hair follicle development (**Fig. 2c**). Specifically, Wnt and Eda signaling pathways were enriched in our study, which had been previously demonstrated to play an important role in mouse hair induction (Chen et al., 2012; Zhang et al., 2009). In addition, Wnt and Notch signaling had been demonstrated to take part in mouse hair differentiation (Lin et al., 2000; Ouji et al., 2008) (Fig. S1). To confirm the expression pattern of the DEGs, we randomly selected 4 genes *(Vcan, Fn1, Tgfbi, Sox9)* to validate their expression patterns using qRT-PCR (**Fig. 3a**). The results were in accordance with the RNA-seq data, suggesting that the expression patterns based on RNA-seq data were reliable.

**Fig 3.**
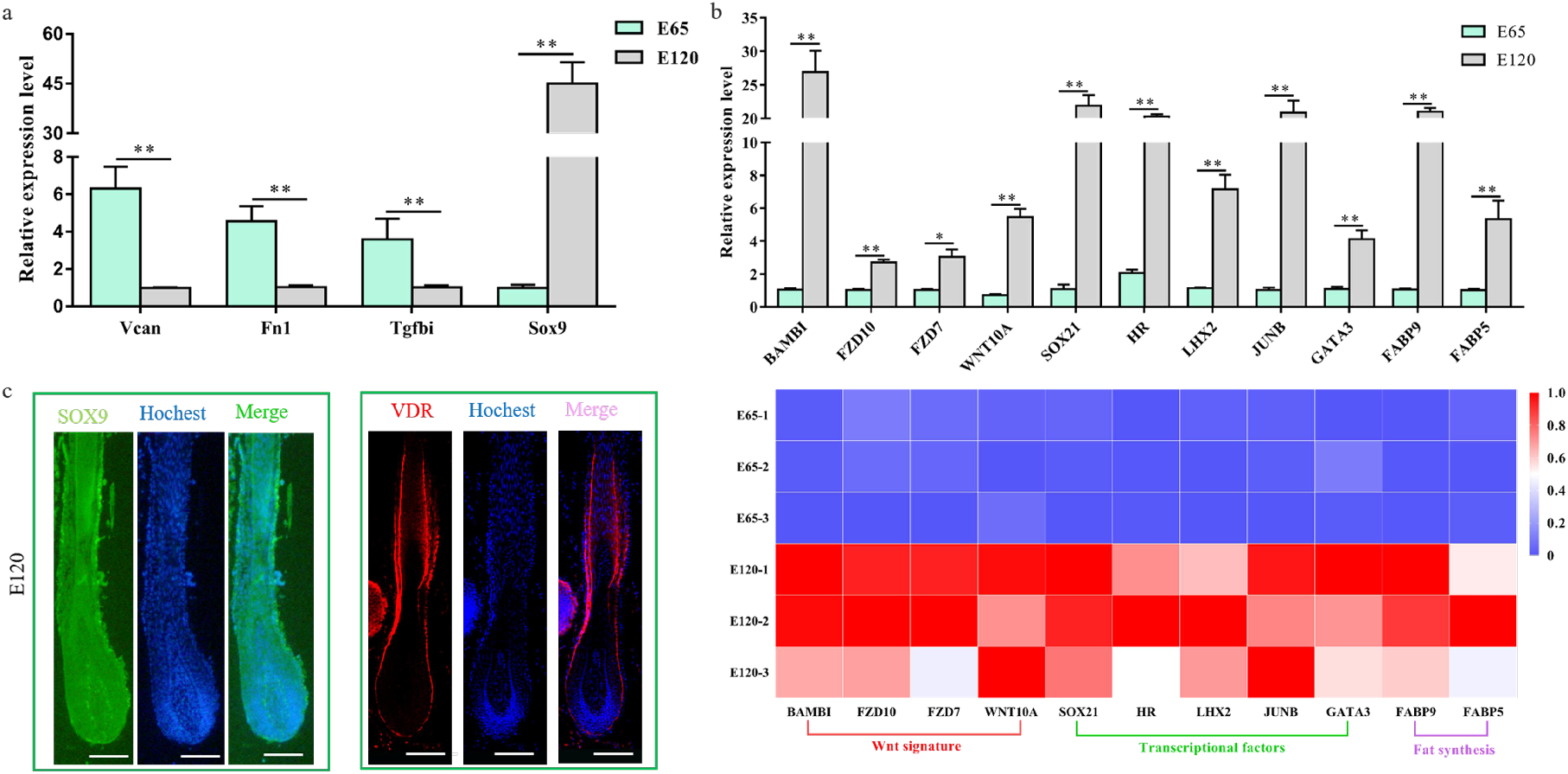
Verification of differentially expressed genes of hair follicle induction and differentiation (**a**) qRT-PCR of four randomly selected genes between E65 and E120 in cashmere goat. (**b**) qRT-PCR confirmed the expression of partial DEGs associated with hair follicle development between E65 and E120 in cashmere goat. And the heatmap was based on the results of qRT-PCR and standardized by Min-max normalization method. (**c**) IF of Sox9 and Vdr at E120 of cashmere goat. Green/Red fluorescence indicated the expression pattern of interest protein. Nucleus was stained with Hoechst in blue. Scale bars, 50 μm.

We revealed that a number of keratin and keratin-associated protein genes were up-regulated or specifically expressed in E 120 (Additional file 1), which was in accordance with the phenotype of hair shaft development in E 120 and that keratin and keratin-associated protein are major structural proteins of hair shaft (Rogers, 2004). Correspondingly, signaling genes belong to Wnt and Notch pathways were up-regulated in E 120, at the same time, transcriptional factors including Hoxc13, Sox9, Sox21, Junb, Lhx2, Vdr, Dlx3 and Gata3 (Dunn et al., 1998; Hwang et al., 2008; Jave-Suarez et al., 2002; Kaufman, 2003; Powell et al., 1992; Törnqvist et al., 2010; Vidal et al., 2005) were up-regulated or specifically expressed in E 120 detected by RNA-seq (Additional file 1), qRT-PCR (**Fig. 3b**) and semi-quantitative RT-PCR (Fig. S2). Furtherly, the expression of Sox9 and Vdr was reconfirmed using Immunofluorescence (IF), the results showed that Sox9 mainly expressed in the outer root sheath and Vdr mainly expressed in the outer root sheath and hair shaft in E 120 (**Fig. 3c**), while not expressed in E 65 (data not shown). These results highlighted the central roles of these transcriptional factors and signals in hair follicle differentiation. Besides, we found several specific genes, which were the critical genes in specific cell types –Pc and DC during hair follicle morphogenesis at E 14.5 in mice (Ahn, 2014; Biggs and Mikkola, 2014; Sennett et al., 2015) were expressed at E 65 of cashmere goat (FPKM >0.5) (Fig. S3), which indicated that these genes may play important roles in hair induction. To further validate the specificity of these genes, we performed IHC validation. The result showed that Edar, Bmp2 and Fgf20 specifically expressed in Pc, while Bmp4 specifically expressed in DC (**Fig. 4a-d**), which suggested that these genes could be the markers for Pc and DC in cashmere goat.

**Fig 4.**
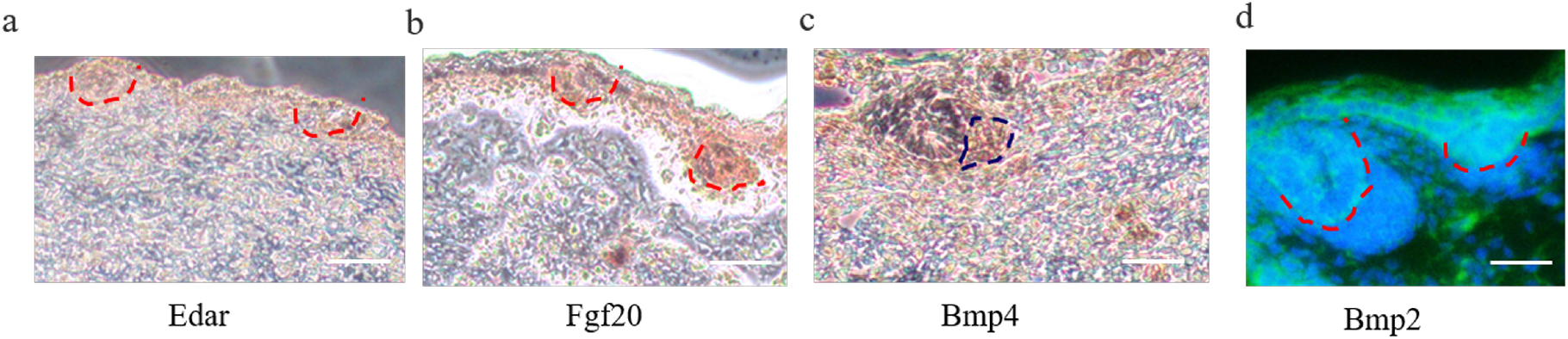
IHC verification of Pc and DC cell-type-specific gene on skin tissues. (a-d) Edar, Bmp2 and Fgf20 specifically expressed in Pc, while Bmp4 specifically expressed in DC. Brown indicates the expression of interest protein. Green fluorescence indicates the expression pattern of Bmp2, nucleus was stained with Hoechst in blue. Red dashed lines indicate epidermal hair follicle placode, blue dashed lines indicate dermal condensate. Scale bars, 50 μm.

### Wnt signal in hair follicle induction and differentiation

From our study and previous studies, Wnt signaling is one of the foremost signaling during hair induction and hair differentiation (Chen et al., 2012; Glover et al., 2017). However, which cell generates the Wnt signal molecules and which cell receives the signal during hair induction is still unclear. β-catenin is stabilized expressed in the nucleus when extracellular Wnt proteins bind to Frizzled receptors and low-density related lipoproteins in the target cell membrane (Tsai et al., 2014). Hence, in our study, we detected the expression of β-catenin using IF to reflect the activated Wnt signal. The result revealed that β-catenin was expressed in epidermal hair follicle placode (**Fig. 5a**), suggesting Wnt signal is activated in epidermal cells during hair induction. Consistent with that, Fzd10, the receptor of Wnt ligands, was also expressed in epidermal hair follicle placode (**Fig. 5c**). Meanwhile, Wnt ligands are lipid-modified extracellular glycoproteins that require the activity of wntless protein (Wls) for secretion (Bänziger et al., 2006). In order to investigate which cell emits the Wnt ligands, we examined the expression of Wls protein by IF on dorsal skin at E 65. Wls protein was detectable in the surface ectoderm as cytoplasmic staining and was enriched in the early developing hair follicle placode rather than dermal cells (**Fig. 5b**). The result suggested that the Wnt signal in hair placode was activated under the control of Wnt ligand from hair placode. At E 120, Wls and β-catenin were expressed in outer root sheath, matrix and hair shaft (**Fig. 5d-e**) which was in accordance with previous studies in mice (Millar et al., 1999), suggesting Wnt signal also play an important role in cashmere goat hair differentiation.

**Fig 5.**
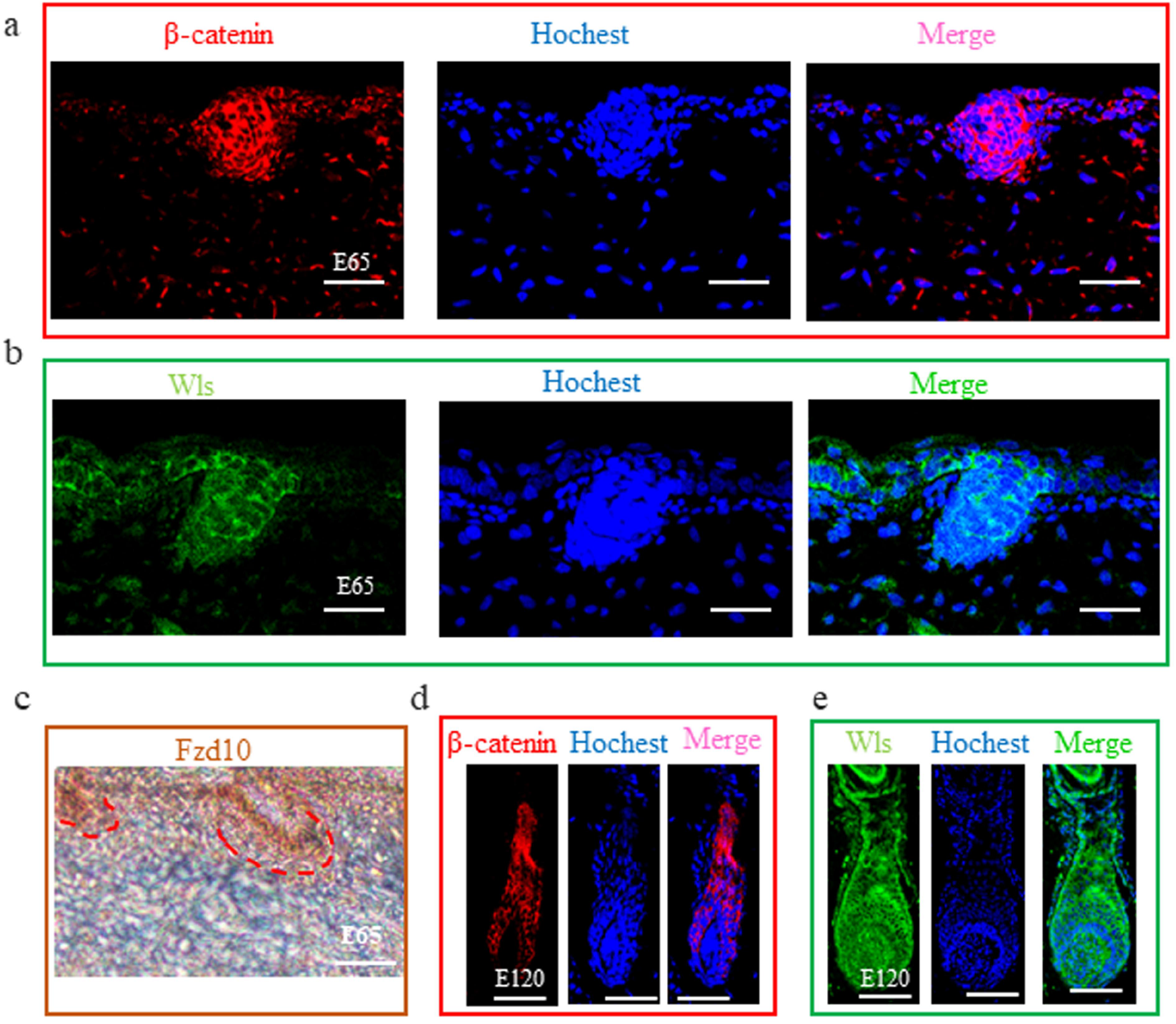
The expression of β-catenin, Wls and Fzd10 at hair follicle induction and differentiation stages were detected by IHC. (**a-c**) The expression of β-catenin, Wls and Fzd10 at E65 stage. (**d-e**) The expression of β-catenin and Wls at E120 stage. Green/Red fluorescence indicated the expression pattern of interest protein. Nucleus was stained with Hoechst in blue. Brown indicates the expression of Fzd10 protein. Red dashed lines indicate epidermal hair follicle placode. Scale bars, 50 μm.

### LncRNA analysis of skin hair follicle development

To investigate whether lncRNA takes part in DNA methylation and plays an important role in hair follicle induction and differentiation, lncRNA transcriptome from RNA-seq was analyzed to define the lncRNA patterns in E 65 and E 120 skin tissues. After rigorous process of selection and coding potential analysis using the software CNCI (https://github.com/www-bioinfo-org/CNCI), CPC (http://cpc.cbi.pku.edu.cn/) and Pfam-scan (http://pfam.xfam.org/), 1407 annotated lncRNAs (Additional file 4) and 13881 novel lncRNAs loci (Additional file 5) including long intergenic non-coding RNA (lincRNAs), intronic lncRNA and anti-sense lncRNAs were identified (**Fig. 6a** and **b**). Compared with protein coding transcripts, lncRNA showed shorter ORF length, transcript length and less exon number (**Fig. 6c**).

**Fig 6.**
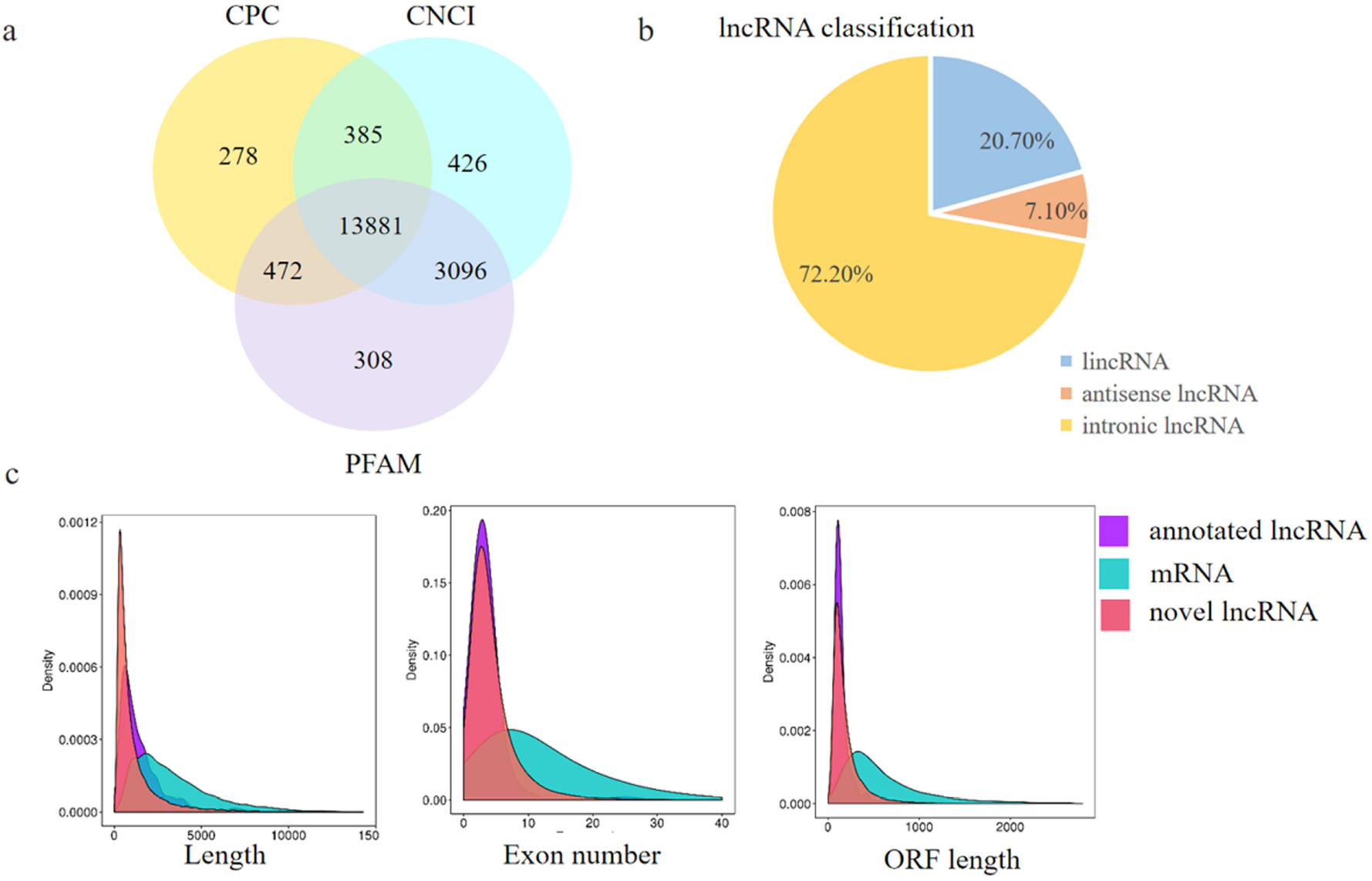
Identification and characterization of lncRNAs in E65 and E120 skin tissues of Capra hircus. (**a**) Screening of the candidate lncRNAs in skin transcriptome by CPC, CNCI and PFAM. (**b**) The classification of lncRNAs. (**c**) Distribution of transcript lengths, exon number and ORF length in the lncRNAs and protein-coding transcripts.

Using edgeR, the differentially expressed lncRNAs (Fold Change ≥2 and P-adjust value ≤0.05) between E 65 and E 120 were screened, resulting in 192 differentially expressed lncRNAs including 45 up-regulated and 147 down-regulated lncRNAs in E 120 compared with E 65 (Fig S4a) (Additional file 6). Meanwhile, a few lncRNAs were specifically expressed at a single developmental stage of hair morphogenesis, such as lnc_006636 showed E 65-specific expression, while lnc_000374, lnc_001937 and lnc_009323 showed E 120-specific expression, indicating that these lncRNAs could regulate cashmere morphogenesis through their spatio-temporal expression. Subsequently, we randomly selected 5 differentially expressed lncRNAs to validate their expression patterns using qRT-PCR. The results were in accordance with the RNA-seq data and showed that lnc_000374 and lnc_002056 specifically expressed at E 120 (**Fig. 7**), suggesting that the expression patterns based on RNA-seq data were reliable.

**Fig 7.**
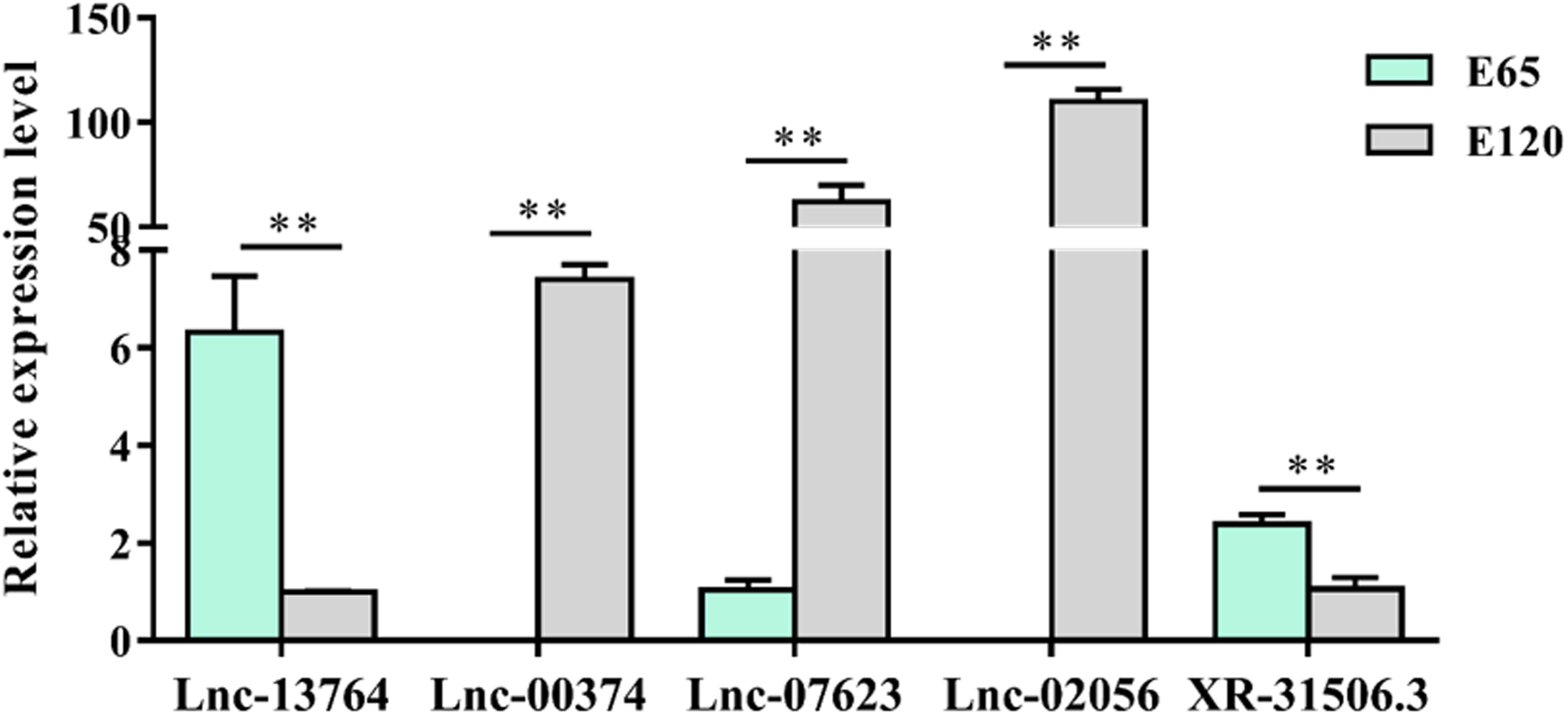
The lncRNA expression patterns in different stages. β-actin was used as reference genes.

To investigate the function of lncRNAs, the potential targets of lncRNAs in *cis* and *trans* were predicted. For the *cis* action of lncRNAs, we searched for protein-coding genes 100 kb upstream and downstream of the lncRNAs. For the *trans* role of lncRNAs in protein-coding genes was examined based on its expression correlation coefficient (Pearson correlation ≥ 0.95 or ≤ −0.95). Subsequently, KEGG analysis was performed on these target genes. As a result, the target genes enriched in hair follicle related signaling pathways including Wnt, Focal adhesion and Ecm receptor pathway (Fig. S4b), indicating lncRNA may participate in hair induction and differentiation through regulating related target genes.

### Genome DNA methylation of hair induction and differentiation during morphogenesis

We found the differential genes between E 65 and E 120, which indicated that hair morphogenesis is the consequence of the spatial and temporal expression of genes. As known, DNA methylation plays a critical role in those genes’ expression (Suzuki and Bird, 2008). However, the regulation mechanism of DNA methylation during hair morphogenesis remains unknown in cashmere goat. Therefore, we detected the DNA methylation at E 65 and E 120 (n=3) skin tissues using WGBS. A total of 195.37 G and 187.09 G raw data were generated for the E 65 and E 120 groups, respectively. An average of 212 million raw reads of WGBS data for the E 65 and E 120 groups were analyzed. Approximately 90.20 % (E 65) and 89.6 % (E 120) of clean reads could be independently mapped to the goat reference genome assembly ARS1 (Table S2 and S3). Any ambiguously mapped and duplicate reads were removed from downstream analysis. Then, the methylation levels of each cytosine were calculated.

An average of 1.78 % and 1.97 % methylated cytosines (mCs) of all genomic C sites in E 65 and E 120 were detected, respectively (Table S4), suggesting the mC level in hair follicle induction stage was higher than that in hair follicle differentiation stage during hair follicle morphogenesis. Methylation in goats was found to exist in three classifications: mCG, mCHH (where H is A, C, or T), and mCHG, in which mCG was the predominant type (>96 %) in both E 65 and E 120 groups. To examine the overall methylation status, methylation levels in different genetic structural regions were determined, including promoters, exons, introns, CpG islands (CGIs) and CGI shores (regions within 2 kb of an island). At the genome-wide scale, the E 65 samples (hair follicle induction stage) exhibited a higher CG methylation status in all regions (**Fig. 8**), which indicated that demethylation took place in E 120 (hair follicle differentiation stage) to ensure the cell lineages. In accordance with that, qRT-PCR showed that Tet3, which intermediates in the process of DNA demethylation as DNA hydroxylases, was expressed higher in E 120 compared with E 65 (Fig. S5). Meanwhile, a marked hypo-methylation was observed in the regions surrounding transcription start site corresponding with the previous studies (Jones, 2012).

**Fig 8.**
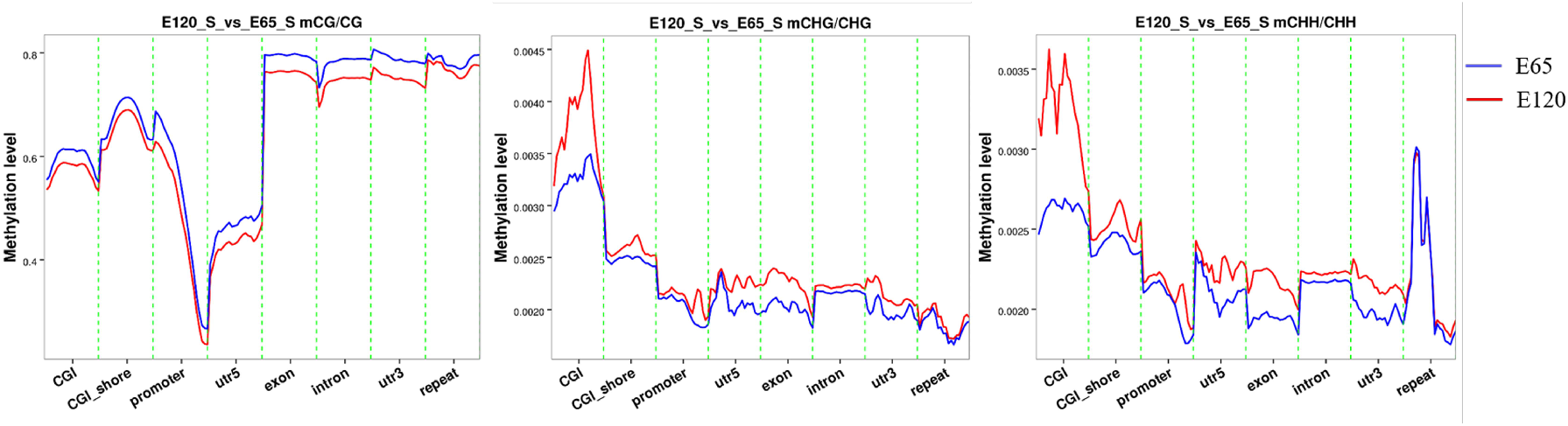
The methylation level in gene different regions for mCG, mCHH and mCHG. At the genome-wide scale, the E65 samples exhibited a higher CG methylation status in all regions. A marked hypo-methylation was observed in the regions surrounding transcription start site.

To identify genomic regions with different levels of methylation between the E 65 and E 120 stages, methylated residues were examined by analyzing sliding windows of 1000 bp in length using DSS. A total of 6899 differentially methylated regions (DMRs) were identified, including 5241 hyper DMRs and 1658 hypo DMRs in E 120 compared with E 65 (Additional file 2). Next, the genes within the DMRs were annotated using the ARS1 assembly. The analysis revealed a total of 3371 genes that were determined to be differentially methylated genes (DMGs) (**Fig. 9a**). To obtain a better mechanistic understanding on the gene regulatory networks controlled by DNA methylation that may be responsible for functional differences during hair induction and differentiation. KEGG analysis revealed that the DMGs were enriched in TGF-β and Focal adhesion signaling pathways (**Fig. 9b**). These results highlighted the central roles of DNA methylation regulation in intercellular crosstalk and signaling transduction during hair follicle induction and differentiation.

**Fig 9.**
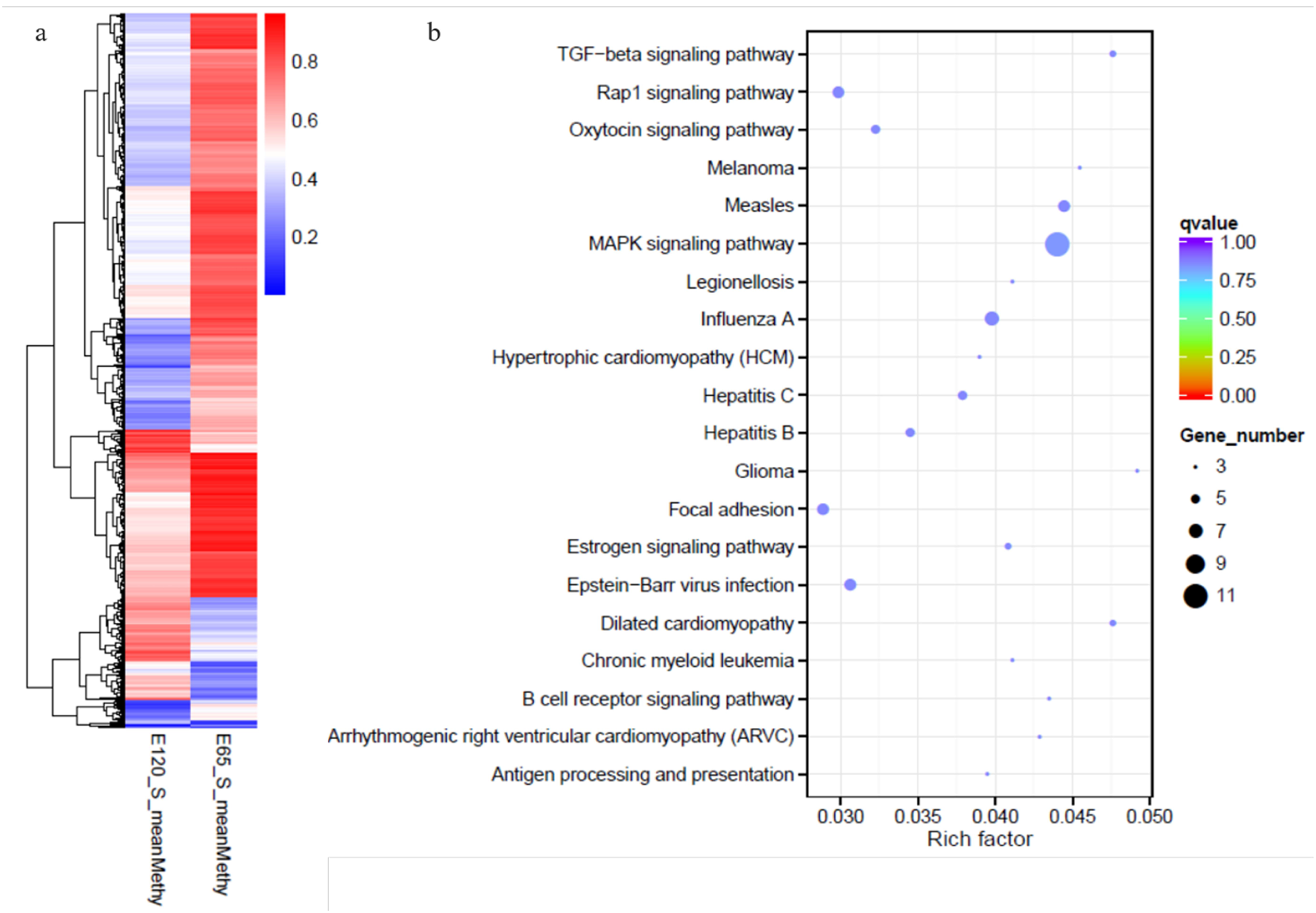
The heat map and KEGG analysis of genes with differential methylation between E65 and E120 (a) The heat map of the genes with differential methylation between E65 and E120. (b) The KEGG analysis of the genes with differential methylation between E65 and E120.DEG-upgenes DMR-Hypogenes DEG-downgenes DMR-Hypergenes

### Integrated analysis of WGBS and mRNA-seq data

To determine the relationship between DNA methylation and gene expression, the integrated analysis of WGBS and RNA-seq data was performed. As a result, we detected 547 hypo-methylation genes with higher expression in E 120 while 282 hyper-methylation genes with lower expression in E 120 compared with E 65 (Additional file 3) (**Fig. 10**). In order to verify the relationship between DNA methylation and gene expression, four genes involved in hair follicle development were selected to reconfirm using BSP-seq and qRT-PCR. The result of BSP-seq was in accordance with that of the WGBS, and the gene expressions were in accordance with the RNA-seq data, in which the genes was repressed by the high DNA methylation (**Fig. 11**).

**Fig 10.**
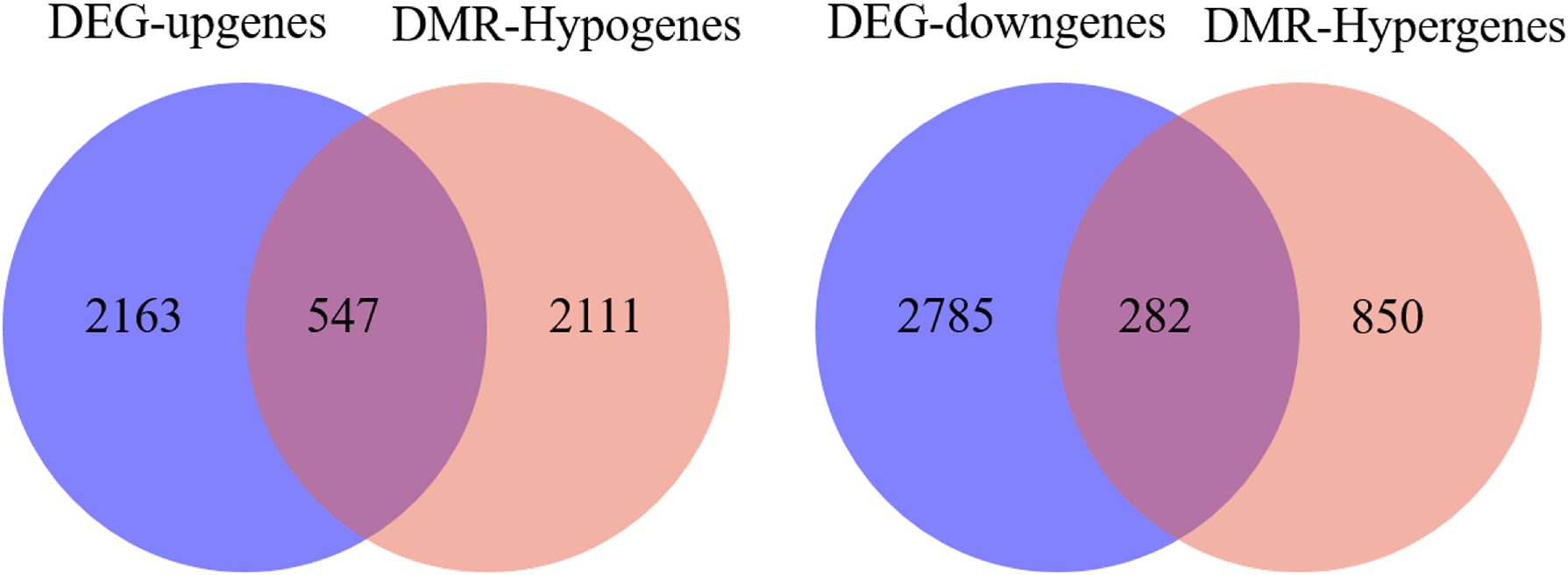
The Venn diagram between the differentially methylated genes and differentially expressed genes between E65 and E120. A. hypo-methylation genes with higher expression B. hyper-methylation genes with lower expression.

**Fig 11.**
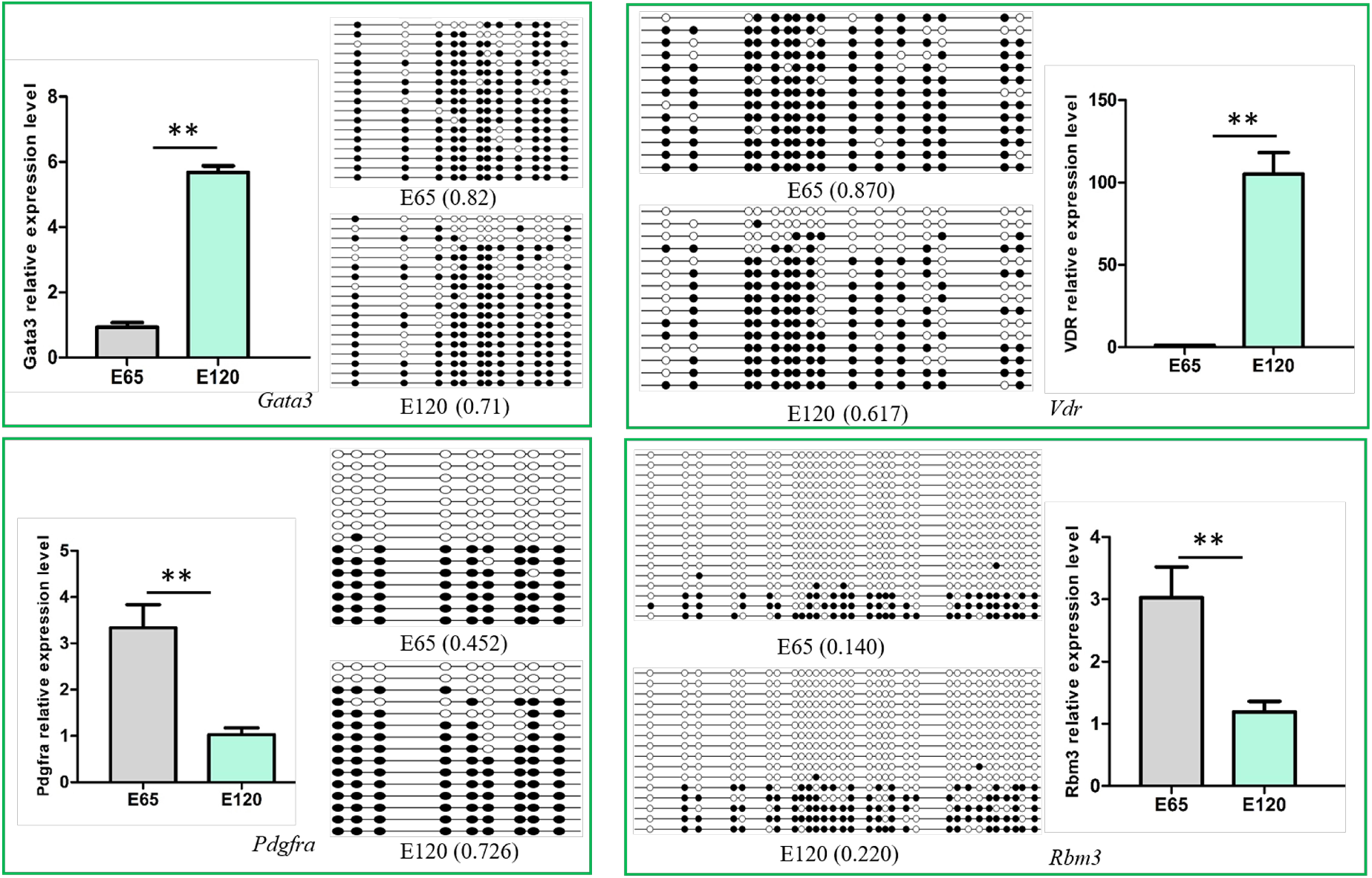
Verification of the differentially methylated genes and their expression. β-actin as a reference gene for quantitative gene expression. The data was expressed as the mean± SE (n=3). ** P<0.01.

It was noteworthy that the transcriptional factor genes associated with hair differentiation including *Gata3, Vdr, Cux1, Tp63* and *Runx1* had low expression with high DNA methylation during hair induction stage in our integrated analysis on WGBS and RNA-seq data. Meanwhile, the signaling genes associated with hair differentiation and development including *NOTCH1, NOTCH3, JAG1, FZD1, SMAD7* and keratin gene *KRT40* had similar expressions and DNA methylation patterns with above transcriptional factor genes (Table 1). The results suggested that DNA methylation play an important role in hair differentiation through regulating associated gene expression. Hair differentiation-related genes did not express in hair induction stage with high methylation, while expressed with hypo-methylation when hair differentiation. Demethylation may occur in hair differentiation to regulate DNA methylation and gene expression.

**Table 1.** The genes associated with hair follicle differentiation under the control of DNA methylation

| Gene | E120<br>FPKM | E65<br>FPKM | log2(fold<br>change) | Pvalue | E120<br>mean<br>Methy | E65 mean<br>Methy | Start | end |
| --- | --- | --- | --- | --- | --- | --- | --- | --- |
| *Tp63* | 48.8 | 5.3 | 3.2 | 0.005 | 0.35 | 0.70 | 77226924 | 77227046 |
| *Vdr* | 19.3 | 0.0 | Inf | 0.000 | 0.73 | 0.91 | 31960999 | 31961087 |
| *Gata3* | 11.1 | 2.1 | 2.376964 | 0.005 | 0.58 | 0.83 | 12388542 | 12388674 |
| *Cux1* | 5.0 | 0.0 | 11.11304 | 0.011 | 0.52 | 0.84 | 35639445 | 35639901 |
| *Runx1* | 3.1 | 1.0 | 1.589259 | 0.002 | 0.65 | 0.19 | 146939482 | 146939629 |
| *Gli3* | 5.1 | 3.1 | 0.716708 | 0.003 | 0.44 | 0.88 | 41410654 | 41411108 |
| *Foxo1* | 11.6 | 1.4 | 3.049298 | 0.000 | 0.40 | 0.72 | 64694258 | 64694508 |
| *Fzd1* | 18.1 | 5.8 | 1.64596 | 0.004 | 0.33 | 0.68 | 111902257 | 111902552 |
| *Notch1* | 18.1 | 6.3 | 1.528654 | 0.003 | 0.71 | 0.86 | 103351558 | 103351699 |
| *Notch3* | 10.6 | 5.0 | 1.091672 | 0.017 | 0.51 | 0.87 | 100777692 | 100777863 |
| *Smad7* | 7.6 | 1.4 | 2.415998 | 0.005 | 0.36 | 0.80 | 48827145 | 48827355 |
| *Jag1* | 33.5 | 8.3 | 2.015405 | 0.001 | 0.33 | 0.79 | 3752377 | 3752803 |
| *Rora* | 23.5 | 4.2 | 2.489794 | 0.003 | 0.55 | 0.87 | 53503152 | 53503499 |
| *Egfr* | 30.8 | 13.0 | 1.237949 | 0.004 | 0.32 | 0.68 | 842607 | 843297 |
| *Fgfr2* | 3.2 | 0.9 | 1.783451 | 0.007 | 0.44 | 0.70 | 10433298 | 10433462 |
| *Krt40* | 28.6 | 0.1 | 8.433506 | 0.000 | 0.39 | 0.79 | 40830782 | 40831215 |
| *Krt14* | 1321.9 | 6.6 | 7.638095 | 0.000 | 0.66 | 0.90 | 41440140 | 41440345 |

### Potential lncRNA that could take part in DNA methylation

Furtherly, in order to investigate the function of lncRNA on gene expression regulation through mediating DNA methylation, integrated analysis of lncRNA, mRNA transcriptome and WGBS were performed. As a result, the potential differential expressed lncRNAs associated with DNA methylation on target genes were revealed (Additional file 7). Such as, lncRNA XR_001918556 may affect the DNA methylation of transcriptional factor gene *Gata3*, lnc-013255 may affect the DNA methylation of transcriptional factor gene *Tp63*, lnc-003786 affect *Fgfr2*, lnc-002056 affect *teneurin-2* which encodes transmembrane proteins. Lnc-007623 may affect the DNA methylation of *Add1* gene, which encode a cytoskeletal protein. The lncRNA expression patterns in different tissues of E120 and skin different stages were show in figure S6. We found lnc-002056, lnc-007623 and lnc-000374 were specifically expressed in skin tissue at E120, corresponding with the hyper DNA methylation of their target genes at E120, which indicated their potential role on DNA methylation regulation.

## Discussion

Mouse pelage hair follicle formation has been divided into nine distinct developmental stages (0-8) for twenty years (Paus et al., 1999). Increasing functional molecules have been identified and characterized for each stage using spontaneous mouse mutants and genetically engineered mice(Nakamura et al., 2013; Saxena et al., 2019; Sundberg et al., 2005). However, there are few reports regarding the machinery underlying cashmere goat hair follicle morphogenesis due to technical difficulties and high costs. Although there are conservative signals in hair follicle development among mammals, however, different physiology and regulation mechanism exists between mouse and cashmere goat. Cashmere is nonmedullated and under the control of seasonal variation of light, which is different from mice (Ge et al., 2018; Mcdonald et al., 1987). Further evidencing the differences is the fact that, *EDAR* gene-targeted cashmere goats showed different phonotypes in hair follicle compared with the targeted mice (Hao et al., 2018; Srivastava et al., 1997). As hair follicle morphogenesis and development determine the yield and quality of cashmere, it is critical to reveal the underlying molecular mechanism. Hence, based on H&E staining results, E 65 and E 120 samples were selected to identify the signals and genes involved in hair induction and differentiation stages.

Hair follicle morphogenesis relies on the interaction between epidermal and dermal cells, ultimately resulting in differentiation of hair shaft, root sheaths, and dermal papilla (Rogers, 2004; Saxena et al., 2019). Corresponding with that, through RNA-seq and bioinformatics analysis, the DEGs were found related to signaling, cell migration and aggregation highlighting the central roles of intercellular crosstalk and dynamic cell rearrangement in hair morphogenesis. Specifically, Wnt signal has been demonstrated play a critical role in hair induction (Andl et al., 2002; Zhang et al., 2008). However, accurate signal transmission between different cells is still unknown during hair induction. Through IF of β-catenin and Wls, we revealed that the Wnt signal in hair placode is activated under the control of Wnt ligand from hair placode. Meanwhile, a number of keratins had a similar expression pattern with some transcriptional factors, which specifically expressed in E 120, suggesting that these transcriptional factors played critical roles in hair follicle differentiation and keratin expression. Furthermore, the signature genes for Pc and DC were found through comparing with related reports on mice (Sennett et al., 2015), the result contributed to illustrating the accurate signal communication between different cells and could be used as markers to isolate specific cells.

During early embryonic development, cells start from a pluripotent state, from which they can differentiate into multiple cell types, and progressively develop a narrower differentiation potential (Wolf, 2007). Their gene-expression programs become more defined and restricted, in which DNA methylation play a critical role in this process (Michael et al., 2007; Ozkul and Galderisi, 2016). Unlike embryonic stem cells, progenitors are restricted to a certain lineage but have the potential to differentiate into distinct terminal cell types upon stimulation. During hair morphogenesis, hair progenitor cells start in a multipotent state, from which they can differentiate into many hair cell types, and progressively develop a narrower potential (Klose and Bird, 2006; Senner, 2011; Wolf, 2007). However, the DNA methylation changes of lineage-committed progenitors to terminally differentiated cells are largely unknown. Recently, researches demonstrated that DNA methylation is a critical cell-intrinsic determinant for astrocyte, muscle satellite cells and mammary epithelial cells differentiation and development (Dirk, 2015). Sen et al. revealed that the dynamic regulation of DNA methylation patterns was indispensable for progenitor maintenance and self-renewal in mammalian somatic tissue. DNMT1 protein was found enriched in undifferentiated cells, where it was required to retain proliferative stamina and suppress differentiation (Sen et al., 2010). Li revealed that DNA methylation played an important role in maintaining hair follicle stem cell homeostasis during its development and regeneration (Li et al., 2012). However, the DNA methylation change during hair morphogenesis is still unknown. In our study, we revealed that the DNA methylation was lower in hair follicle differentiation compared with hair follicle induction stage. Furtherly, hair follicle differentiation genes including transcriptional factors and signaling genes were methylated in hair induction stage but were subsequently de-methylated during differentiation. The result suggested that DNA methylation patterns are required for hair induction and differentiation. Correspondingly, Bock revealed that DNA methylation changes were locus specific and overlapped with lineage-associated transcription factors and their binding sites, which played an important role during in vivo differentiation of adult stem cells (Bock et al., 2012) and that demethylation events were frequently linked to brain specific gene activation upon terminal neuronal differentiation (Guo et al., 2014).

Another related report in Shanbei cashmere goat, Li et al. previously revealed that DNA methylation had little effect on gene expression when telogen-to-anagen transition in adult Shanbei White cashmere goat (Li et al., 2018). Combined with the above researches, the DNA methylation patterns from oocyte to adult including hair follicle morphogenesis and cycling were described in Shanbei White cashmere goat (**Fig. 12**). DNA methylation was higher in adult than embryonic period and had little change when telogen-to-anagen transition in adult Shanbei White Cashmere goat. The darker color represents the higher DNA methylation. The results will enrich the regulatory network of hair morphogenesis.

**Fig 12.**
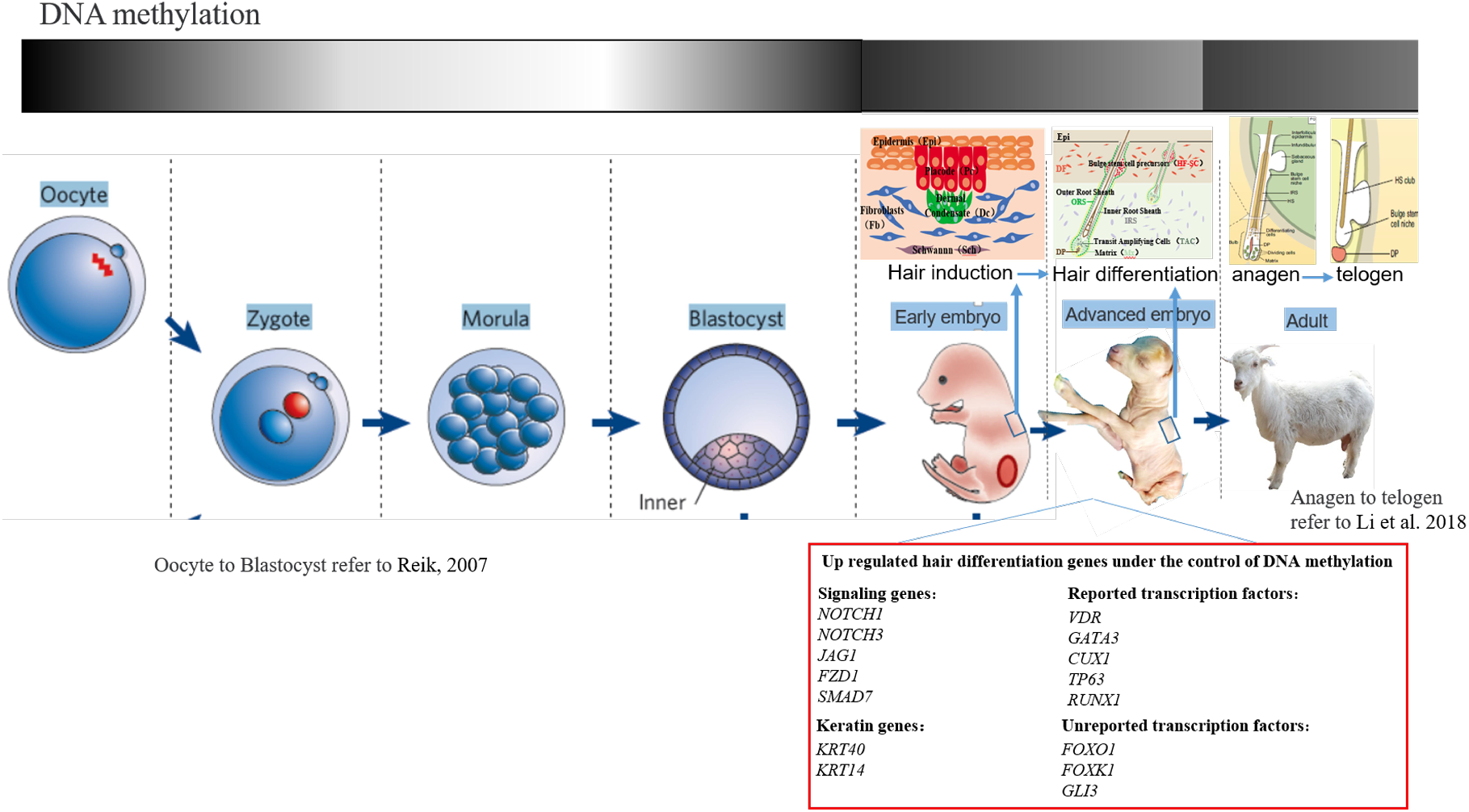
The dynamic changes of DNA methylation from oocyte to adult in cashmere goat

Above, we revealed that locus specific DNA methylation changes played a critical role during hair morphogenesis. However, both DNA methyltransferases and polycomb repressive complexes lack sequence-specific DNA-binding motifs. Increasing evidence indicates that many lncRNAs contain DNA-binding motifs that can bind to DNA by forming RNA:DNA triplexes and recruit chromatin-binding factors to specific genomic sites to methylate DNA and chromatin (Han and Chang, 2015; Sun et al., 2016). Besides, lncRNA have been associated with important cellular processes such as X-chromosome inactivation, imprinting and maintenance of pluripotency, lineage commitment and apoptosis (Engreitz et al., 2013; Mohammad et al., 2012; St et al., 2015). However, the function of lncRNA in hair morphogenesis is still unknown. In our study, we revealed lncRNA may have a function through targeting hair follicle related signals and genes. Furtherly, potential lncRNA involved in DNA methylation was revealed. However, the specific function of lncRNA need to be further studied. The results provide a potential regulatory mechanism mediated by lncRNA during hair morphogenesis.

## Conclusions

In this study, the critical signals and genes were revealed during hair follicle morphogenesis in cashmere goat. In this process, differentiation genes were governed by DNA methylation, resulting in repressed expression in hair follicle induction stage and high expression in hair follicle differentiation stage. Furtherly, potential lncRNAs associated with DNA methylation on target gene were revealed. This study would enrich the regulatory network and molecular mechanisms in hair morphogenesis.

## Supplementary Materials

Fig. S1: The heatmaps of DEGs associated with signaling pathways related to hair follicle development, Fig. S2: Semi-quantitative RT-PCR confirmed the expression of partial DEGs associated with hair follicle development between E65 and E120 in cashmere goat, Fig. S3: The potential cell-type-specific markers during hair induction and differentiation in Shanbei White Cashmere goat, Fig. S4: Differentially expressed lncRNAs and their KEGG analysis in cashmere goat skin between E65 and E120 during hair morphogenesis, Fig. S5: Tet3 was expressed higher in E120 compared with E65, Fig. S6: The lncRNA expression patterns in different tissues of E120 and skin different stages, Table S1: Primer list for qRT-PCR, Table S2: Data statistics of WGBS at E65 and E120 of cashmere goat, Table S3: The quality control of WGBS data at E65 and E120 of cashmere goat, Table S4 The statistics of methylation in genome scale at E65 and E120 of cashmere goat.

## Author Contributions

Conceptualization, Xin Wang and Shanhe Wang; software, Shanhe Wang; validation, Shanhe Wang, Fang Li, Yuelang Zhang and Yujie Zheng.; resources, Lei Qu and Jinwang Liu; data curation, Xin Wang and Fang Li.; writing—original draft preparation, Shanhe Wang; visualization, Shanhe Wang and Wei Ge; supervision, Xin Wang; project administration, Xin Wang and Lei Qu; funding acquisition, Xin Wang.

## Acknowledgements

This research was funded by the National Natural Science Foundation of China (No.31772573).

## Accession Numbers

All the RNA-seq data and WGBS data sets supporting the results of this article have been submitted to the National Center for Biotechnology Information (NCBI) Gene Expression Omnibus (GEO). mRNA-seq data: SAMN13669153, SAMN13669154, SAMN13669155, SAMN13669156, SAMN13669157, SAMN13669158. WGBS-seq data: SAMN13679866, SAMN13679867, SAMN13679868, SAMN13679869, SAMN13679870, SAMN13679871.

## Appendix

Additional file 1: The differential expressed genes between E65 and E120 stages during hair follicle morphogenesis in cashmere goat (xlsx, 4406 KB)

Additional file 2: Differentially methylated regions between E65 and E120 during hair morphogenesis in cashmere goat (xlsx, 899 KB)

Additional file 3: The DEGs negatively correlated with DNA methylation between E65 and E120 of cashmere goat (xlsx, 119 KB)

Additional file 4: The sequences of annotated lncRNAs in E65 and E120 skin of cashmere goat (xlsx, 2352 KB)

Additional file 5: The sequences of novel lncRNAs in E65 and E120 skin of cashmere goat (xlsx, 17465 KB)

Additional file 6: The differential expressed lncRNAs between E65 and E120 stages during hair follicle morphogenesis in cashmere goat (xlsx, 69 KB)

Additional file 7: The potential differentially expressed lncRNAs associated with DNA methylation on target genes (xlsx, 631 KB)

